# Musical interaction is influenced by underlying predictive models and musical expertise

**DOI:** 10.1101/440271

**Authors:** Ole A. Heggli, Ivana Konvalinka, Morten L. Kringelbach, Peter Vuust

## Abstract

Musical interaction is a unique model for understanding humans’ ability to align goals, intentions, and actions, which also allows for the manipulation of participants’ internal predictive models of upcoming events. Here we used polyrhythms to construct two joint finger tapping tasks that even when rhythmically dissimilar resulted in equal inter-tap intervals (ITIs). Thus, behaviourally a *dyad* of two musicians tap isochronously at the same rate, yet with their own distinct rhythmical context model (RCM). We recruited 22 highly skilled musicians (in 11 dyads) and contrasted the effect of having a shared versus non-shared RCM on dyads’ synchronization behaviour. As expected, tapping synchronization was significantly worse at the start of trials with non-shared models compared to trials with a shared model. However, the musicians were able to quickly recover when holding dissimilar predictive models. We characterised the directionality in the tapping behaviour of the dyads and found patterns mostly of mutual adaptation. Yet, in a subset of dyads primarily consisting of drummers, we found significantly different synchronization patterns, suggesting that instrument expertise can significantly affect synchronization strategies. Overall, this demonstrates that holding different predictive models impacts synchronization in musicians performing joint finger tapping.

**Public significance statement:** This study shows that when a pair of musicians thinks differently about a rhythm they play together, their performance is worse. However, they are able to recover back to normal performance levels after a few taps for which they use different strategies. Interestingly, we find that the strategies used by drummers may be different from other musicians.

## Introduction

Successful interpersonal interaction and communication is dependent on shared predictive models^1–4^. We understand others’ actions by inferring their goals, beliefs, and attitudes^5,6^. Recently, musical paradigms have proven very useful for studying interpersonal interaction mechanisms, since precise predictive models can be easily experimentally manipulated (e.g tonality or meter) and quantified using interaction dynamics, which occur at a timescale of milliseconds^7^. Here we used a minimalistic musical paradigm to investigate the interactions between a dyad of two musicians having the same or different top-down predictions.

Humans are highly adept at coordinating movements with one another and do so often without noticing, which again influences social behaviour. Hence, interpersonal coordination may be a human predisposition, evident in behaviours such as the tendency towards synchronised walking^8^, or how dyads in rocking chairs attempt to synchronize even when the natural frequencies of their chairs are incongruent^9^. This tendency to synchronize movements is also visible in body movement in more complex interactions, such as the bodily synchronization during joke telling^10^. Synchronization in these tasks appears to be spontaneous, seemingly without conscious effort. In contrast, musical performance requires the ability to synchronize movements on-the-fly between performers, and is a case of intentional interpersonal coordination. Music consists of temporally related sounds, where movements must be precisely coordinated both within and between performers^11,12^. Thus, in music, interpersonal coordination is a result of intended joint action rather than a spontaneous occurrence or basal mimicry.

Much of interpersonal coordination research – and joint action more broadly – has been concerned with understanding the mechanisms that enable people to successfully coordinate with one another^13^ – specifically, the interplay between the top-down predictive and bottom-up reactive mechanisms. It is now well established that we make continuous predictions about the sensory consequences of our own actions, allowing us to attenuate perceptions of predicted sensations and amplify salience of externally caused ones^14^. Prediction hence plays a major role in joint action, embedded within a common coding framework of action and perception, scaling across the what, when, and where of another’s actions - i.e. *what* another’s intention to act is, *when* they will act temporally, and *where* in the common space they will act ^15^. Joint action may thus best be understood within a predictive coding framework^6,16–19^.

Within this framework, perception and action are governed by top-down processing^17^. From the viewpoint of joint action, this means that assumptions about the interaction are translated into predictions of perceptual input – one person’s action output becomes another’s perceptual input, and vice-versa^2^. These predictions are then compared with the perceived input. If they are found to be incoherent, then bottom-up reactive signals serve as prediction errors towards the purpose of forming a revised assumption. In this sense, our interaction with others and the world as a whole relies on forming, testing, and revising predictive models^6^.

In music, the top-down predictions are formed not only by interactions, but also by an internal model of certain basic constituents of the music itself such as melody, harmony, and rhythm^20^. Manipulating these internal models for musical interaction provides a unique opportunity for experimentally manipulating and measuring the influence of the individual top-down predictions on joint action. In the present study we asked two participants to tap isochronously together, i.e. producing a rhythm based on holding one of two distinct meters (here called rhythmical context models, RCM).

Meter serves as a common denominator for most types of music, and especially Western music^21^. The meter is a hierarchical framework consisting of evenly spaced and differentially accented beats, providing to each metric position a timing and a metrical weight. Examples of different meters are the rhythmical framework underlying waltz (3/4) or march (2/4 or 4/4). The metrical weights are thought to linearly correspond to the strength of the expectation towards events occurring at these time points^22^. In other words, the more metrically salient a position is in the hierarchy, the stronger the expectation that events will occur at this metrical position. The ability to perceive meter is fundamental not only to music, but also to language perception, and in motor control^23^. The ability to detect a regular pulse in auditory stimuli may be innate, as shown in EEG-studies demonstrating that infants (2-3 days old) are able to detect deviations in beat^24^.

Hence, rhythm perception is conceptualized as the interplay between the brain’s anticipatory structuring of music and what is heard, forming the RCM^25^. This model is not necessarily stationary and immutable, but rather constantly updated through the process of musical interaction. Examples of this would be the waxing and waning in tempo found in classical music, or in the organic change to double-time in a jazz performance^26^. In fact, since the meter is a mental model, it is possible to perceive a piece of music from the point of view of two different meters. In the case of a polyrhythm, two different perceptual interpretations of the exact same auditory stimulus may be equally plausible^27^.

A classic example of a polyrhythm is the 3-against-4 rhythmical pattern. It may be experienced by playing, for example on the drums, at the same time three equally spaced beats with one hand and four equally spaced beats with the other, so that the periods of both patterns add up at the same period of time. For such rhythmical patterns, it is possible to perceive the meter as either a duple meter (formally 4/4) with the three-beat pattern as the counter-metric pattern (Figure 1a), or, alternatively, as a triple meter (formally 3/4) with the four-beat pattern as a counter-metric pattern (Figure 1b)^28,29^. The rhythmic organization of the two interpretations in Figure 1 is exactly the same, that is, the cross-rhythmic relationship between the two streams within each pattern is identical. These two experiences of the same polyrhythm (albeit with inverted instrumentation) are phenomenologically different and is thus analogous to ambiguous images such as Rubin’s vase, which can be seen either as a vase on black background, or as faces on white background^28^. As with Rubin’s vase^30^, cross-rhythm in music can sometimes cause perceptual shifts in which the metric model is reinterpreted as one (triple) or the other (duple).

**Figure 1.**
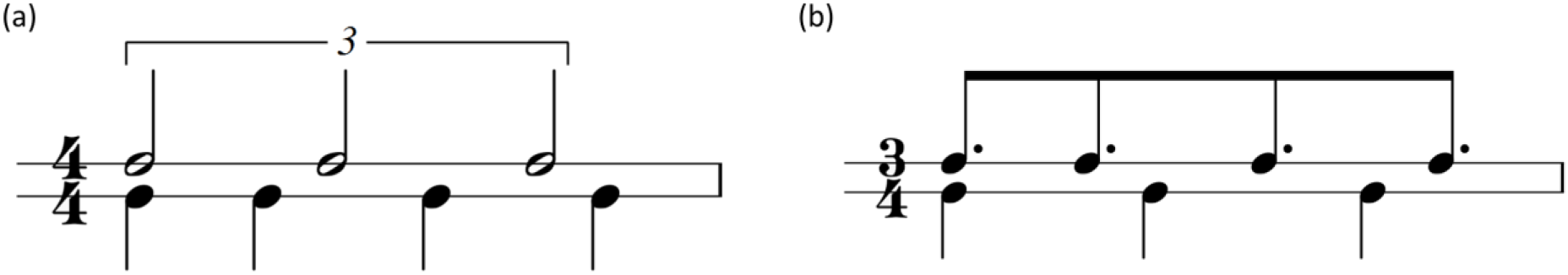
3-against-4 pattern. In (a) the meter is 4/4 with the three-beat pattern as the counter-metric pattern. In (b) the meter is 3/4, with the four-beat pattern as the counter-metric pattern.

Rubin’s vase is a visual example of ambiguous sensory input which is perceived different depending on the underlying mental models. Even though such fundamentally different perception probably often goes unnoticed during everyday interaction they occasionally surface, such as in the Internet phenomenon of 2015 of naming the colour of a dress (search for the widely used #theDress)^31^. However, interactions between perception when using different mental models and subsequent behaviour have never been studied experimentally. In the visual example above, the mental model switches involuntarily between the two percepts. For polyrhythm such as 3 against four, however, it is possible to make skilled musicians maintain one stable meter (3/4 or 4/4) throughout a musical excerpt.

Here, we investigated the behavioural effects of holding one of two RCMs when producing and hearing the same rhythmic isochronous pattern (Figure 2a). In more detail, we designed a joint finger-tapping paradigm which had identical rhythms, i.e. motor output, irrespective of whether the underlying RCMs were shared or non-shared. A shared RCM occurred when both participants were tapping the same rhythm, and hence having the same top-down predictive model (see Figure 2c). In contrast, when one participant was tapping the polyrhythm and the other a straight rhythm, the dyad has two conflicting predictive models resulting in a non-shared RCM. Crucially, these rhythms can be constructed so that they result in an equal motor output, which allowed us to hide the fact that the dyads were actually tapping different rhythms. Hence, we were able to quantify the effects of the internal predictive model on synchronization within the dyads. Previous research has emphasized the necessity of shared predictive models of successful interactions, and we therefore hypothesized that a non-shared RCM would result in decreased synchronization measures^19^. We expected this effect to occur due to discrepancy between the perceived stimuli and the internal predictive model. In particular, the polyrhythm used in the experiment has a bar length of 1.5 seconds whereas the straight rhythm has a bar length of 2 seconds. Thus, any rhythmical adjustment done on the timescale of the bar will be different between the two.

**Figure 2.**
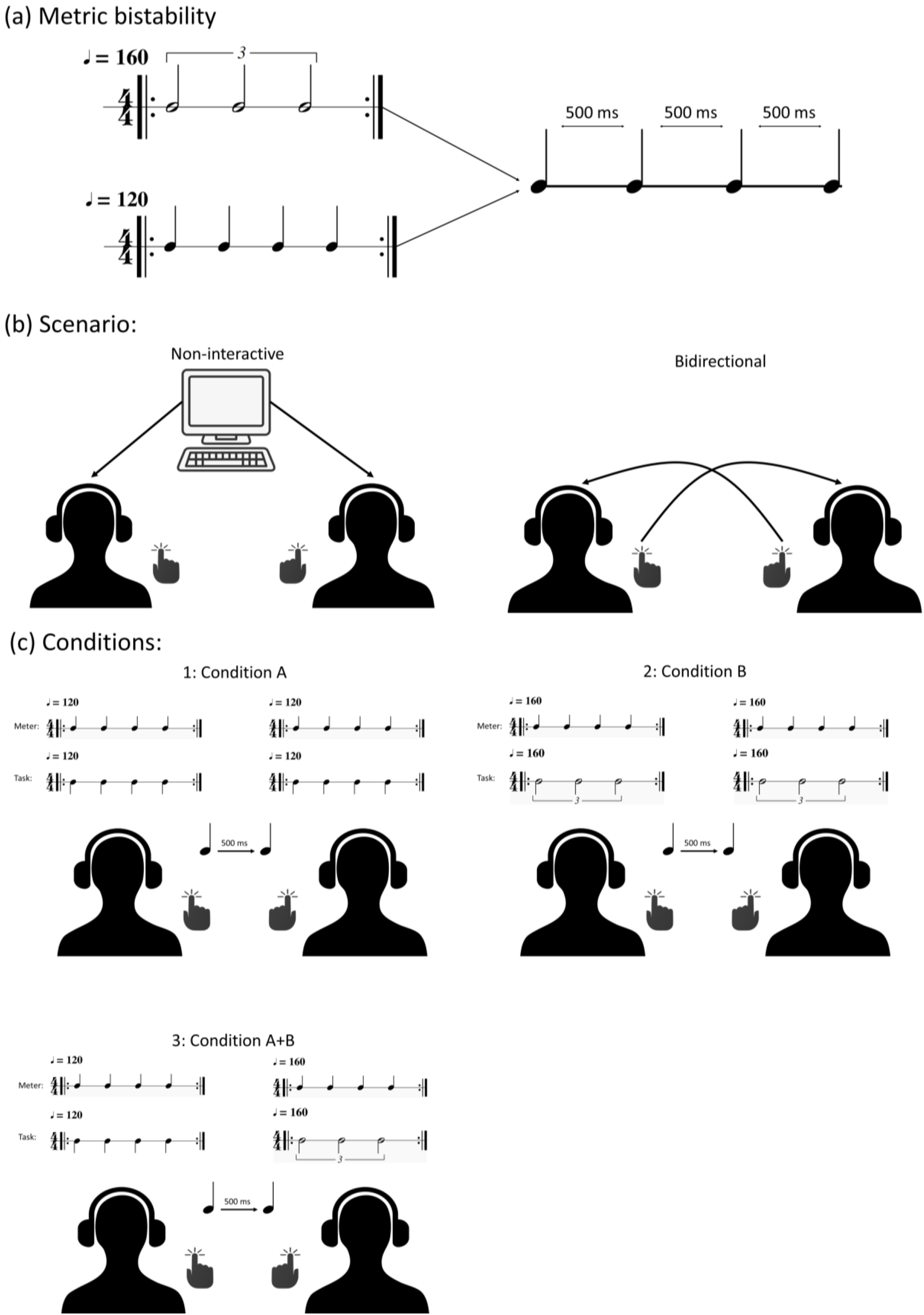
A: An illustration of how an isochronous rhythmic sequence with a 500 ms ITI can be resolved into two distinct rhythms – a 3-against-4 polyrhythm at 160 BPM, and a 4/4 straight rhythm at 120 BPM. B: The two scenarios in our experiment. Participants either tapped along with a computer metronome – the non-interactive condition; or in a bidirectionally coupled state. C: An overview of the conditions, showing both the meter and the task. Note that condition A+B is balanced, so that both member 1 and member 2 performs the two distinct rhythms an equal amount of times.

Further, based on earlier studies, we hypothesized that a leading-following strategy would be more prominent in the non-shared RCM condition^2,4^ (for an overview of synchronization strategies, see Figure 3). While the underlying mechanisms of synchronization strategies remains a topic of discussion, recent research suggests that mutual adaptation is the most efficient strategy and is premediated by a merging of self-other representation^19^. In joint finger tapping tasks, this would entail considering the auditory feedback created by the other as linked to one’s own action. In contrast, the leading-following strategy necessitates a consistent categorizing of the auditory feedback as belonging to the other person. In the non-shared RCM condition, we hypothesized that differences in tapping would force the members into considering the auditory feedback as dissociated from their own tapping. From this, one of the members, presumably the one with the most confidence in their own RCM, would then gravitate towards a leading role.

**Figure 3.**
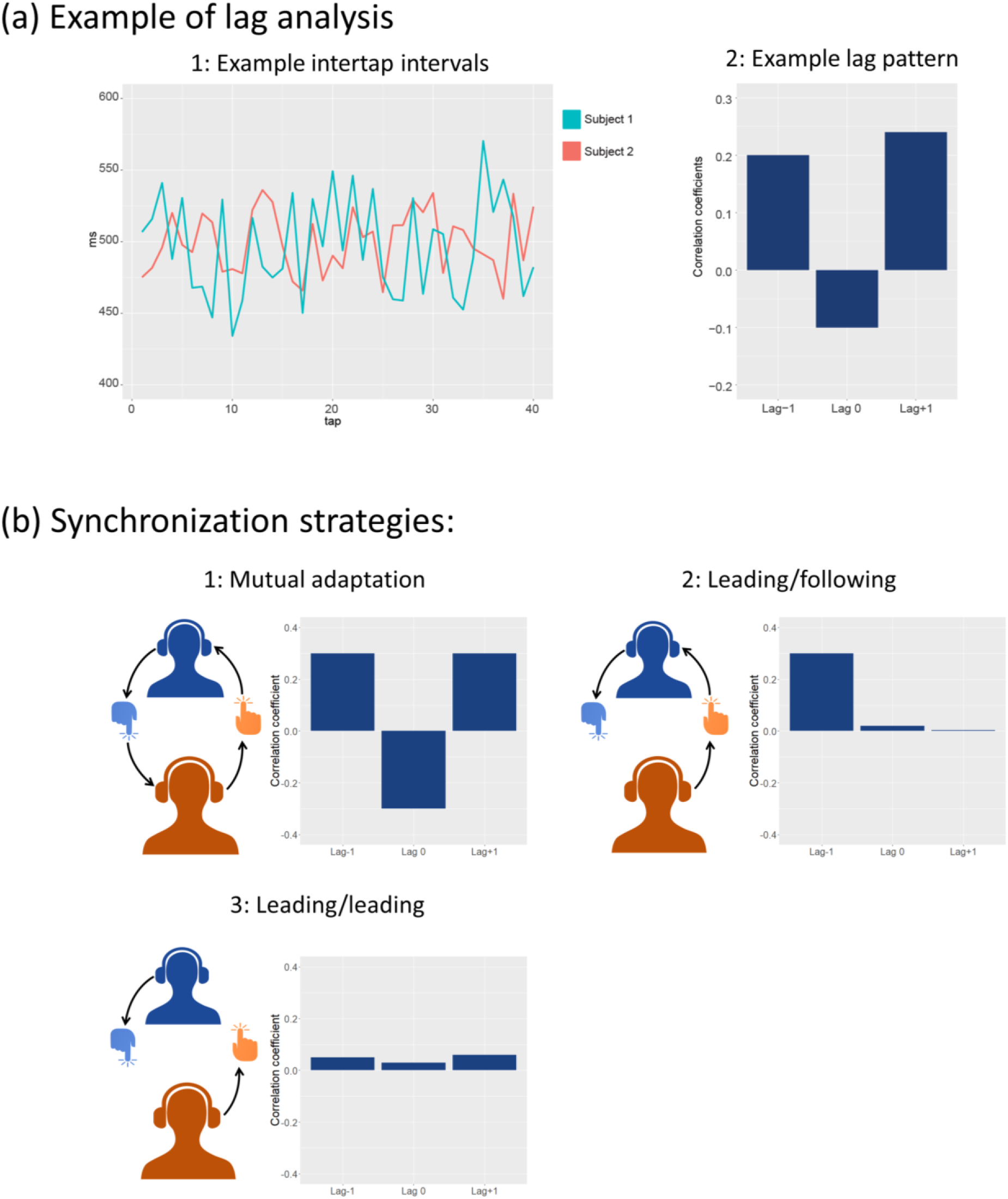
A: An example of the lag analysis. Here, simulated data is shown exhibiting a mutual adaptation dynamic. In the first graph the ITI of a simulated sequence is shown. Cross-correlation is performed on the ITI timeseries, giving correlation coefficients for lag −1, 0 and +1. B: An illustration of different synchronization strategies and their corresponding lag patterns. Mutual adaptation occurs when a perception-action loop is formed between the two members of a dyad, such that they equally weigh the incoming auditory stimuli from the other member, and their own model of the task. Leading-following depends on one of the dyad members to attenuate the information coming from the other member, here illustrated by the missing connection between the blue member’s tapping and the orange member’s headphone. As such, the leader puts more confidence in their own internal model. In the case of leading-leading, as observed in our experiment, both participants exhibit leading behaviour. Here, they both discard information from the other member and rather relies on their own model.

## Methods

### Ethics statement

This study was conducted at the Center for Music in the Brain in the Department of Clinical Medicine at Aarhus University, and ethical approval were therefore governed by the Central Denmark Region Committees on Health Research Ethics. The committee found that the study was not considered a health research study according to the Act on Research Ethics Review of Health Research Projects (Act 593 of July 14^th^ 2011, section 14.1 and 14.2), and did therefore not require ethical approval (reference number 87/2016). Nonetheless, the experiment was performed in accordance with Aarhus University’s policy for responsible conduct of research and the Declaration of Helsinki. Participants signed consent forms and was informed that their participation was entirely voluntarily, and that they could exit the experiment at any time without any penalty to the payment for participation. In addition, participants were debriefed and given contact information to the researchers performing the experiment. The collected data were anonymized, and no national identification number was stored.

### Paradigm

As we were interested in the interpersonal effects of the RCM, we decided to employ a joint finger tapping paradigm with bidirectional coupling – participant 1 received auditory feedback of participant 2’s taps, and simultaneously participant 2 received auditory feedback of participant 1’s taps. In addition, we added a non-interactive scenario where the participants tapped along with a computer metronome to serve as a baseline condition (Figure 2b). Here, they did not receive any auditory feedback from their tapping. To constrain the amount of shared information within the dyads in the bidirectional condition we restricted auditory feedback from the tapping to only contain a transient sound of equal amplitude and length independent of tapping strength. This way the only information communicated between the pairs were their willingness to adapt, through adjusting their ITI.

### Participants

In order to have participants that could reliably perform the 3-against-4 polyrhythm we recruited highly skilled musicians with normal hearing and no known sensorimotor nor neurological disorders. Participants were recruited predominantly from the Aarhus region, Denmark. A total of 30 paid volunteers participated. They were paired depending on the time slots they signed up for, for a total of 15 pairs with both same and mixed gender. Out of these, 2 dyads were discarded due to their inability to reliably perform the polyrhythm, and one dyad were discarded due to figuring out the hidden condition. The remaining participants self-reported their musical abilities primarily as professional (n=14) and semi-professional (n=9), with one participant self-reporting as amateur. The participant’s mean age was 23.2 years (SD= 2.8), and 2 were female. Their expertise as measured by the mean of years of formal musical training and years of actively playing ranged from 5 to 24 years with a mean of 13.6 years (SD=4.6). Most musicians had a percussion instrument (reported as Drums, Classical Percussion, Percussion, Cajon) as their primary instrument (n=15), with the rest playing harmonic or melodic instruments such as the Piano, Guitar, Saxophone and Trombone.

### Materials and apparatus

Two Arturia Beatstep MIDI controllers were used as tapping devices. MIDI from both devices were recorded on the computer running the experiment, using an M-Audio MIDI-to-USB converter. In order to reduce latency between a participant’s tap and the resulting sound, we used two overclocked Teensy 3.0 microcontrollers, each with an SGTL5000-based audio board. These were connected to the gate voltage output on the MIDI controllers. Using a custom sound generating script, this resulted in tap-to-sound latencies of under 1 ms. The paradigm was programmed in Python, using PsychoPy^32^ and Pyo^33^, presented using a computer running Windows XP, with two monitors – one for each participant.

### Stimuli

Each trial was initiated by one bar of an isochronous and unaccented metronome sound. The metronome stimuli were made with a freely available click sound. For rhythm A, the metronome was four beats at 120 BPM. For rhythm B, the metronome was four beats at 160 BPM. In the bidirectional scenario the metronome stopped after one bar. In the non-interactive scenario, the metronome continued throughout the length of the trial. In condition A+B the metronome for rhythm B started 500 ms later than the metronome for rhythm A, so that the participants would start tapping at the same time.

### Task and procedure

Upon arrival, the members of each pair were given information on voluntarily participating in a research project, received task instructions, and signed consent forms. They were told that they would be participating in an experiment wherein we would be looking at the differences between playing a polyrhythm, and a straight rhythm. As in previous experiments their main objective was to “synchronize with the sound/metronome, and try to maintain tempo”^34^. Participants were given examples of the two tasks they would be performing - a 120 BPM 4/4 simple rhythm (condition A) and the triplet of a faster 160 BPM 3-against-4 polyrhythm (condition B), both resulting in an inter-tap interval (ITI) of 500 ms. They were told there would be two scenarios - a non-interactive computer scenario wherein they would be tapping along with a computer metronome, and an interactive bidirectional scenario wherein they would be tapping along with their partner. In the bidirectional scenario the participants’ auditory feedback was the tapping of their partner, so that dyad member 1 heard member 2’s tapping, and vice versa. In the non-interactive scenario, the participants only heard the computer metronome.

The participants were instructed that a metronome would always start the task with one bar of clicks, and images and text on the screens would inform them of which rhythm to play. Further, they were told to tap with their index finger on a marked pad on the MIDI interface, and to sit as still as possible with their eyes open looking at a fixation cross during the actual tapping. Auditory stimuli were delivered using ER-2 insert earphones (Cortech Solutions), which provided an external noise reduction of >30 dB. Sound levels were set at a comfortable level for each participant. Participants were placed in the same room. While the dimensions of the room made it unfeasible to position the participants back-to-back, they were positioned such that no visual contact was possible without removing the earphones.

For the participants, there were four different tasks – non-interactive: playing a 4/4 simple rhythm (A) referred to as condition A, or the triplet in a 3-against-4 polyrhythm (B) referred to as condition B; and bidirectional: playing rhythm A, or B. However, unbeknownst to the participants there were trials wherein one participant played rhythm A, and the other rhythm B (and vice-versa). This hidden condition, containing the non-shared RCM, is referred to as condition A+B (see Figure 2c). Since we did not want the participants to know that they would be performing different rhythms at the same time, we informed them that they would always tap the same. As their goal ITI would remain the same in all conditions, this was technically correct. An overview of the conditions can be found in Figure 2c. Each scenario had 100 trials, with 50 trials for each rhythm, for a total of 200 trials. This results in 25 trials per scenario for each possible combination of rhythms. Each trial lasted 12 seconds, with the first 2 seconds consisting of the metronome, and the rest of tapping. Example trials can be seen in Figure 5a.

### Data analyses

The paradigm software produced files containing note-on and -off timestamps, which were pre-processed in MATLAB^35^. Only the tapping onset times were analysed, and missing data were dealt with by pair-wise deletion of the onset time. One pair of participants were excluded at this stage of the analysis due to excessive missing data (more than 25% trials with more than 25% missing taps). From the onset times we calculated the inter-tap interval (ITI), the time between two successive taps. Five outcome measures were calculated – ITIs, synchronization indices (SI), cross-correlations, signed asynchrony, and tapping variability as measured by the standard deviation of the asynchronies (SD_asy_). To quantify stability in synchronization behaviour we also calculated the trajectory of the SI during the trial by calculating the average SI at four timepoints consisting of four equal subdivisions of the taps in each trial, so that timepoint 1 represents SI in the first quarter of each trials, timepoint 2 represents the next quarter of each trial, etc. From MATLAB the data was exported to R for statistical analyses^36^. Averages are reported as mean with standard deviation in cases where a normal distribution exists, and median with interquartile range if the distributions did not pass the Shapiro-Wilk test.

To measure how well participants synchronised their tapping to either the metronome or each other, we used synchronization indices (SI). These are calculated based on the variance of relative phase between two signals, giving a unitless number ranging from 0 to 1, based on the following formula^37^:

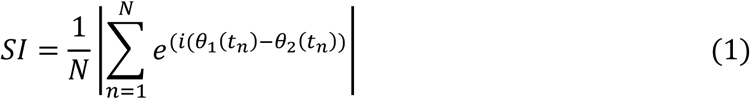

where N represents the number of taps in each trial, and ϴ_1_ and ϴ_2_ are the phases of each dyad member (or computer metronome and participant in the noninteractive scenario). An SI of 1 represents perfect synchronization, and 0 no synchronization. For the non-interactive scenario, we calculated the SI with respect to the metronome signal for each participant, for each of the two rhythms (A and B). The resulting SI were then averaged per participant per rhythm and compared using a Wilcoxon rank sum test. For the bidirectional scenario the SI were calculated between the pair’s time series, indicating how well the participants synchronized to each other. To classify the effect of condition on synchrony while accounting for dyadic differences we used a linear mixed effects model (LMM) with the dyads as the random factor and condition as fixed effect^38^. A rank transformation was applied to the SI, and p-values were obtained by likelihood ratio tests of the model with a null-model retaining only the random factor.

Asynchrony in the bidirectional scenario was calculated as the difference between tap timing of partners in milliseconds. It is signed based on which person is the point of reference, so that when calculated with dyad member 1 being the reference, a positive asynchrony means that member 1’s tap precedes member 2’s tap, and a negative asynchrony the opposite. We calculated the mean asynchrony per trial. The standard deviation of the asynchronies (SD_asy_), was calculated by taking the standard deviation of the absolute asynchronies per trial, and gives an indication of the variability of the finger tapping. A high SD_asy_ indicates that the participants are adjusting their taps over a wide range, whereas a low SD_asy_ indicates a more stable and narrow adjustment range. We used the same LMM approach as described earlier to analyse both the signed mean asynchrony and the tapping variability (SD_asy_), with dyads as random factor, and condition as fixed effect.

To assess directionality in the interaction between the participants we calculated cross correlations at lag −1, 0, and +1 between each dyad member’s tapping time series (Figure 3a). The relation between these coefficients gives an indication as to how the dyad interacts. A perfectly synchronized trial with no variation in tempo would give high correlation at all lags, whereas a trial with high synchrony, but with some variation in tempo, would produce the highest correlation at lag 0. If a leader-follower dynamic is present, a positive correlation at either lag −1 or lag +1 occurs, depending on which participant is the follower (more adaptive one). This occurs due to one member, the follower, lagging one tap behind the leader. Mutual adaptation, wherein both members exhibit following behaviour leads to positive correlation at both lag −1 and lag +1, and negative correlation at lag 0^4^. This lag pattern occurs due to both members constantly adjusting their ITI in opposite directions on a tap-by-tap basis causing the negative correlation coefficient at lag 0. Hence, they are both more correlated to the previous tap of the other dyad member, resulting in an oscillation around an optimal ITI and the corresponding positive correlation at lag −1 and lag +1 (for an overview, see Figure 3).

A challenge in correlation analysis of timeseries such as these is the assumption of stationarity. One way of addressing this issue is to divide the data into overlapping windows, which comes at the cost of making data interpretation harder, due to the increasing risk of spurious correlations^39^. As our participants consisted of highly trained musicians, and our data showed highly stable ITIs, we decided to use conventional cross correlation over the entire trial length. To account for dyadic differences, we performed a two-way MANOVA with condition and dyad as dependent variables on the Fisher Z-transformed lag coefficients.

## Results

### Synchronization indices

A likelihood ratio test revealed a significant effect of condition on synchronization in the bidirectional scenario (χ^2^ (2)=13.03, *p*=0.0015). We performed Bonferroni corrected post-hoc test, using the multcom package in R ^40^. Here, we found that the SI was significantly lowered by approximately −0.023 (*p*=0.0016) in condition B compared to condition A, and by approximately −0.021 (*p*=0.0123) in condition A+B compared to A (see Figure 4). No significant difference between condition B and condition A+B were found, nor between the rhythms in the non-interactive scenario. In the trajectory analysis we performed three separate one-way ANOVAs of the rank-transformed SI over the four timepoints, for each of the condition, and a significant effect was only found in condition A+B (*F(3, 2068)*=4.767, Bonferroni corrected *p*=0.0077) (see Figure 5b). A Tukey HSD test showed that the SI significantly increased between timepoint 1 and 3 (*p*=0.008) and between timepoint 1 and 4 (*p*=0.005). While the change in SI between timepoint 1 and 2 appears to suggest an increase, it only approached significance (*p*=0.067). This means that the synchronization levels increased over time in condition A+B.

**Figure 4.**
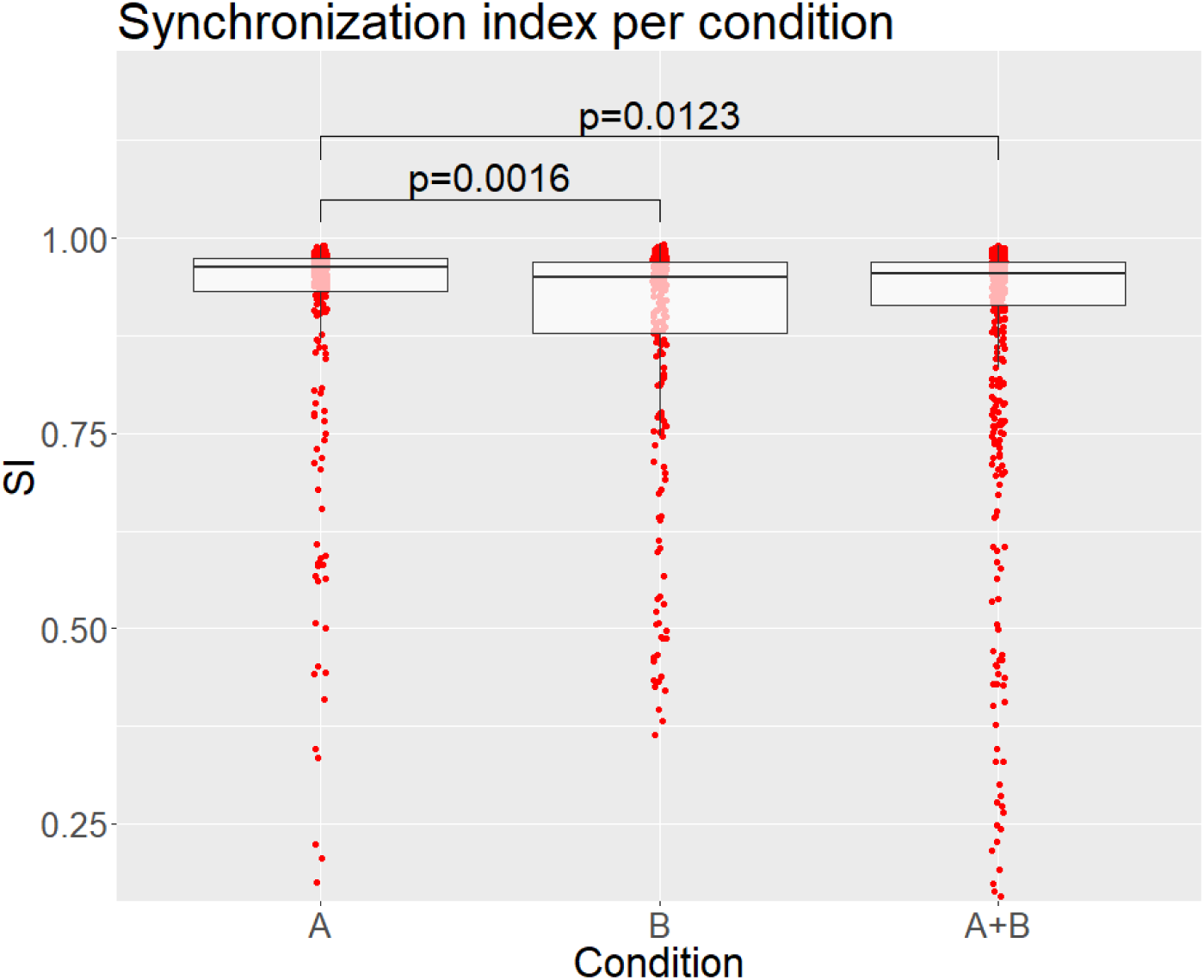
Synchronization per condition in the bidirectional scenario. Individual data points are shown as red dots, and the boxes represent the first and third quartiles, with the median shown as a dark line. P-values are obtained from Bonferroni corrected post-hoc test.

### Tapping variability, SD_asy_

A highly significant effect of condition on SD_asy_ was found using a likelihood ratio test (χ^2^ (2)=16.5, *p*=0.0003). The results mirror the findings from the synchronization index, summarized in table 1. Due to this apparent similarity, we performed a correlation to determine their relationship. A Kendall’s tau-b correlation showed a strong negative correlation between SD_asy_ and SI (tau-b=.496, *p*<0.0001).

**Table 1.** Overview of synchronization measure in the bidirectionally coupled scenario. Values are reported as median and interquartile range, due to non-normal distribution of data.

|  | Condition A | Condition B | Condition A+B |
| --- | --- | --- | --- |
| **SI** | 0.962 (IQR=0.932, 0.973) | 0.949 (IQR=0.877, 0.969) | 0.955 (IQR=0.914, 0.969) |
| **ITI** | 497.3 ms (IQR=490.7, 506.7) | 497.6 ms (IQR=490.1, 517.1) | 497.3 (IQR=490.7, 508.2) |
| **SD<sub>asy</sub>** | 29.6 ms (IQR=25.2, 41.1) | 33.8 ms (IQR=27.7, 45.1) | 32 ms (IQR=26.6, 42.8) |

### Cross-correlations

The cross-correlations of the dyads’ ITI were computed separately and averaged for each condition and for each dyad. A MANOVA using Pillai’s test statistic resulted in no effect of condition (Pillai’s trace=0.317, *F(6, 42)*=1.32, *p*=0.2710), but a significant effect of dyad (Pillai’s trace=1.668, *F(33, 66)*=2.5, *p*=0.0008). An FDR-corrected post-hoc test showed that dyad had a significant effect on all lags (Lag-1 *p*=0.0441, Lag 0 *p*<0.0001, Lag+1 *p*=0.0381), indicating a high between-dyad variability in their tapping strategy.

### Inter-tap intervals

A likelihood ratio test showed significant differences in the participants’ ITIs between the non-interactive and bidirectional scenario. (χ^2^(1)=5.43, *p*=0.0198). In the non-interactive scenario, the median ITI for both rhythms was 500.24 ms (IQR=498.9, 503.1), decreasing to 497.34 ms (IQR=490.7, 510.1) in the bidirectional scenario. Three example trials can be seen in Figure 5a. No significant difference between conditions in either scenario was found.

**Figure 5.**
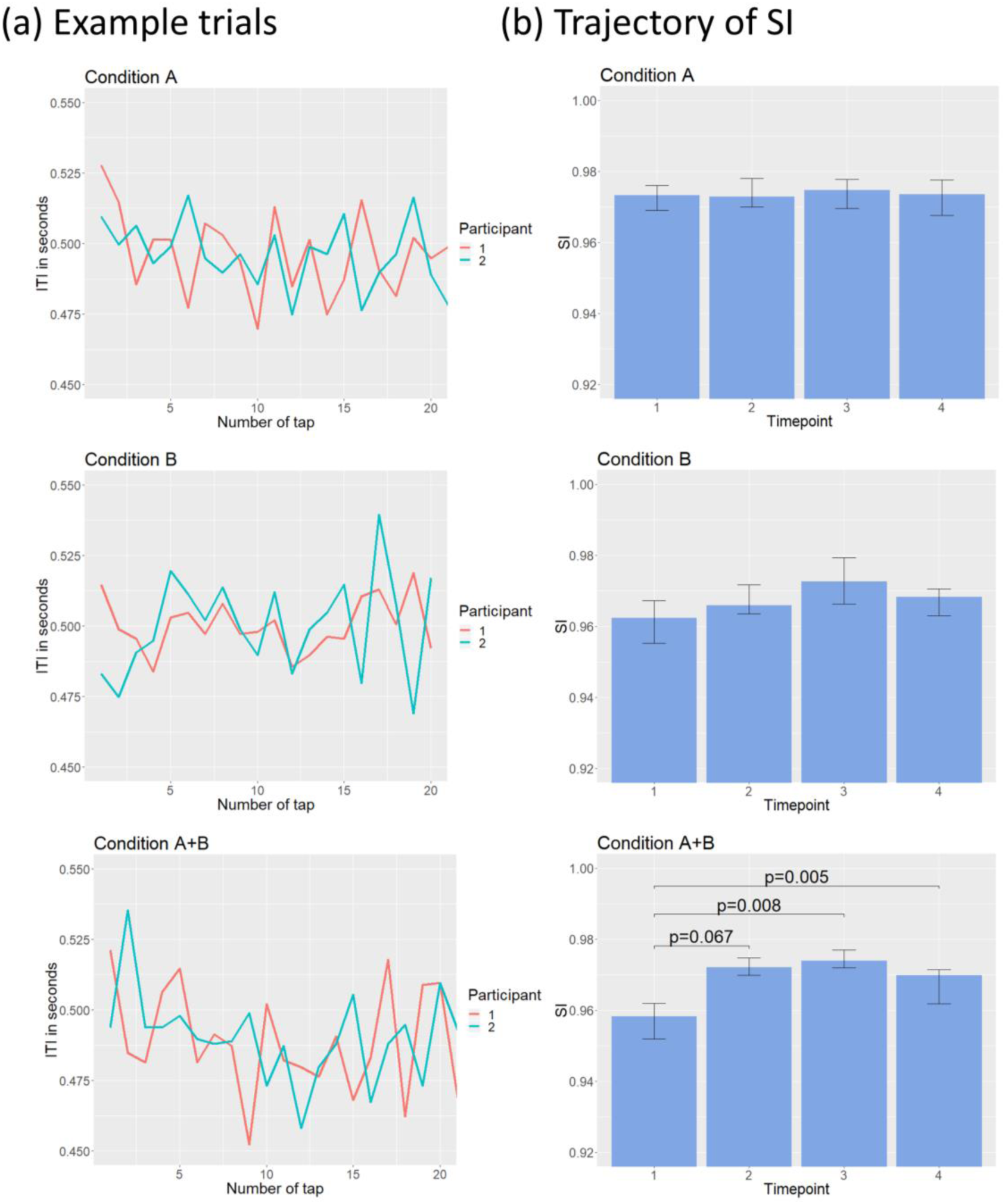
In (a) representative trials from one dyad are plotted as the intertap interval in seconds over the number of taps, for the three conditions. In (b) we show the trajectory of the SI over four timepoints. The timepoints represent the median SI over four equal subdivisions of the taps for each trial. Error bars are bootstrapped 95% confidence intervals. P-values are obtained from a Tukey’s HSD test.

### Asynchrony

Overall, the participants exhibited a low mean asynchrony in the bidirectional scenario of only −3.64 ms (SD=30.8). A likelihood ratio test on the absolute asynchronies only gave a close to significant effect of condition on asynchrony (χ^2^ (2)=4.87, *p*=0.0876).

### Inter-dyad differences in tapping strategy

In order to explore the between-dyad variability in the cross-correlations, we clustered the data by dyad with the lag data as input using Ward’s clustering method (Figure 6a)^41^. A similarity profile analysis using the R-package SIMPROF classified three clusters as significantly different at α<0.05^42^. Visually inspecting the averaged lag coefficients shows that cluster 2 and cluster 3 both exhibit patterns resembling mutual adaptation, whereas cluster 1 shows little to no apparent pattern in its correlation coefficients (Figure 6b).

**Figure 6.**
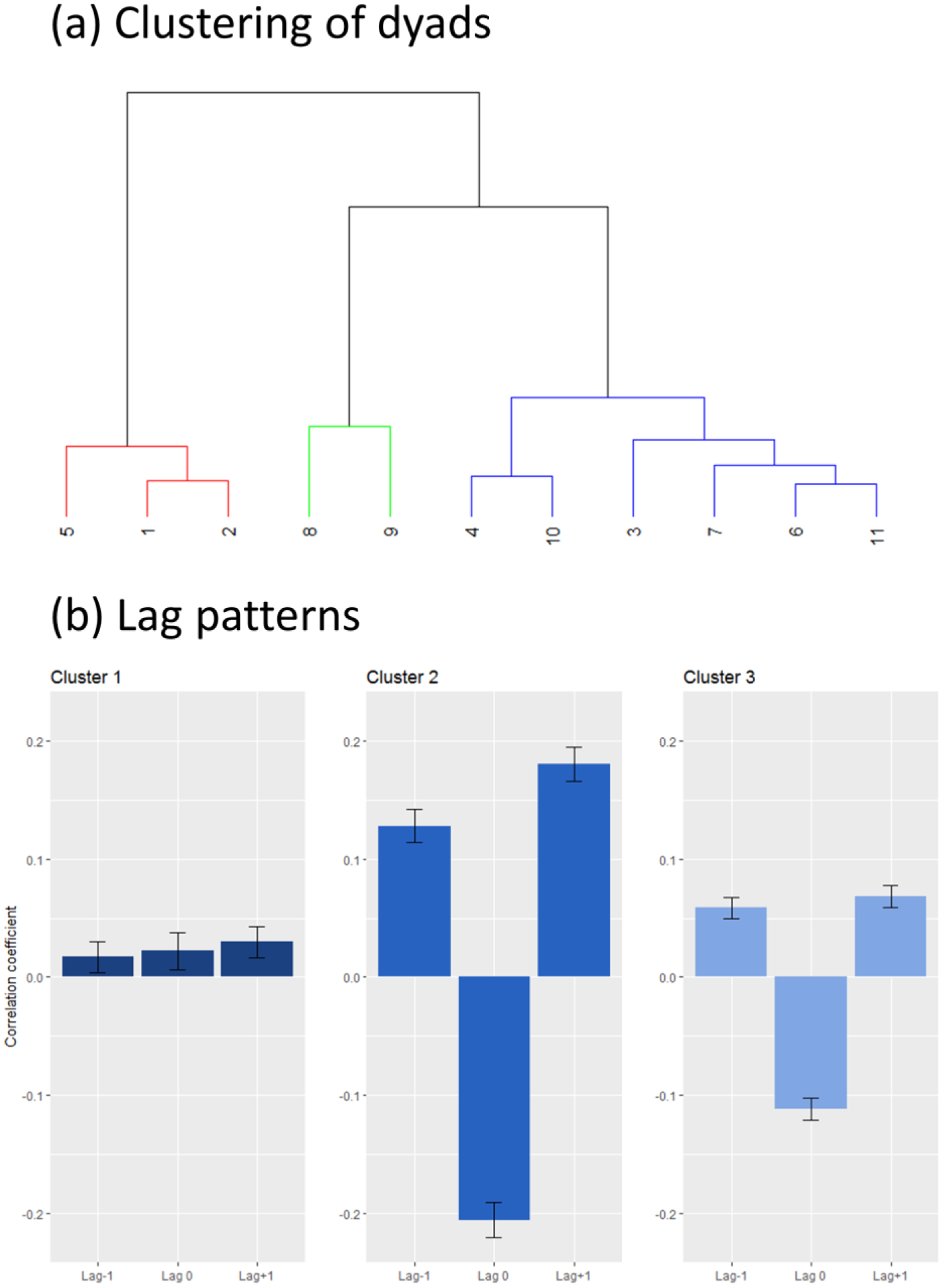
Results from clustering the cross-correlations. A dendrogram of the resulting clusters is shown in A, with colour indicating significant differences at α<0.05. In B, the averaged correlation coefficients for lag −1, lag 0, and lag +1 is shown for each of the clusters, with standard error of the mean error bars. Clusters 2 and 3 can be described as mutual adaptation. Cluster 1 is an example with no clear pattern of interaction, that may come from both participants attempting to lead, causing a leading-leading pattern. Interestingly, this was found primarily in dyads with drummers.

To further classify these clusters, we looked at participants’ primary instruments, categorizing them as 1: drums & percussion (D/P), 2: harmonic & melodic (H/M). We also considered the combinations of primary instruments within a dyad by pairing them in three levels according to whether they consisted of two D/P participants, two H/M participants, or one of each. Further, we classified their experience defined as the mean of years of formal musical training and years of actively playing per participant. We compared these values between the clusters, in addition to the synchronization index and tapping variability. Only instrument and dyad pairing yielded a statistically significant difference (see table 2).

**Table 2.** Characteristics of clusters. For instrument and dyad pairing, we performed a Fisher’s exact test. For experience, SI and SDasy, a one-way ANOVA were performed. We report the median and interquartile range for data that was not normally distributed, and mean and standard deviation for normally distributed data

|  | Cluster 1 | Cluster 2 | Cluster 3 | Statistical value |
| --- | --- | --- | --- | --- |
| **Instrument** | D/P=5 | D/P=0 | D/P=8 | $p = 0.033$ |
|  | H/M=1 | H/M=4 | H/M=4 |  |
| **Dyad pairing** | D/P+D/P=2 | D/P+D/P=0 | D/P+D/P=2 | $p = 0.048$ |
|  | D/P+H/M=1 | D/P+H/M=0 | D/P+H/M=4 |  |
|  | H/M+H/M=0 | H/M+H/M=2 | H/M+H/M=0 |  |
| **Experience** | 12.62 (SD=2.43) | 14.31 (SD=3.1) | 13.13 (SD=5.8) | $F(2,19)=0.155 \ p=0.857$ |
| **SI** | 0.911<br>(IQR=0.676, 0.966) | 0.9579<br>(IQR=0.934, 0.979) | 0.96<br>(IQR=0.937, 0.973) | $F(2,8)=1.17 \ p=0.358$ |
| **SD<sub>asy</sub>** | 12.6 ms, SD=14.4 | 4.1 ms, SD=2.9 | 4.8 ms, SD=7.2 | $F(2,8)=2.715 \ p=0.126$ |

## Discussion

In cases of a non-shared RCM, participants performed significantly better at the end of the trials than at the start. This pattern was not seen in either of the conditions with an equal RCM. This indicates that the internal predictive models impact synchronization when performing joint finger tapping, even though the intended motor outputs in the different conditions are the same. In addition, we identified a subgroup of our participants exhibiting an interaction pattern not reported in coupled interactions before – with very low correlation coefficients across lags −1, 0, and +1, as if they did not interact much with each other, while exhibiting stable tapping behaviour. The subgroup consisted primarily of drummers paired with drummers. Earlier studies primarily report two synchronization strategies in coupled conditions, namely mutual adaptation and leading-following^2,4,34^. We observed a third possible strategy we term leading-leading, wherein synchronization occurs without any apparent dyadic interaction.

Spontaneous synchronization is thought to occur due to a merging of self-other representations, wherein an action-perception loop is created such that one person’s action output becomes another’s perceptual input, and vice-versa^4,19,43^. However, explicit synchronization as encountered in our study may be a more complex phenomenon. Musical rhythmic interaction relies on both maintaining the tempo and synchronizing with other performers, and thus participants may choose to weight these two aspects differently. In our study, we did not find any differences in inter-tap intervals between the bidirectional conditions. The participants performed at a median ITI of 497.34 ms, with an IQR of 490.7 ms to 510.1 ms. This is consistent with research on the just-noticeable differences in tempo sensitivity, which has shown a detection threshold of around 2% difference in sequences at 500 ms^44^, and suggests that the RCM did not significantly influence our participants’ ability to maintain tempo.

While tempo remained relatively stationary, we found differences in synchronization measures. The dyads in our experiment exhibited an increase in SI in the trials with a non-shared RCM. This effect was significant when comparing the first 25% of taps in a trial with the third and fourth 25% of taps. While there was a trend towards an increase in SI in the second 25% of taps compared to the first 25%, this did not reach statistical significance. This effect was not found in neither condition with a shared RCM. A possible explanation here is that the participants start out by keeping their two distinct RCMs, but after recognizing that synchronization is impaired, they abandon their individual RCM and default to a following behaviour. This then leads to an interaction that, as a whole, does not exhibit a large negative effect on synchronization measures. When looking at the entire trials we found that condition A+B was in fact in between condition A and condition B in synchronization measures. The dyads achieved an average SI of 0.955 in condition A+B, as compared to 0.962 in condition A and 0.949 in condition B. Hence, the musicians in our study appear to be resilient to the effects of holding a non-shared RCM.

A plausible explanation of this observed behaviour is found in the brain’s tendency for computational optimization. Viewed from the framework of predictive coding under the free energy principle, the brain constantly attempts to optimize its use of energy by minimizing the error between its predictions and the sensory input^45^. Thus, when humans are interacting, spontaneous synchronization may occur as a result of individuals minimizing the differences in self/other representations^19^. In our study, the decreased SI seen at the start of the non-shared RCM trials may represent the participants individually searching for a magnitude of variability wherein they are able to minimize the difference between self and other representation. Our data then suggests that explicit synchronization such as rhythmic joint action may share many traits with spontaneous synchronization.

Previous studies on joint finger tapping in a bidirectionally coupled setting predominantly report mutual adaptation as measured by cross-correlations^4,34^. Similar patterns of interaction are also found in experiments studying, for instance, imitative hand movements^46^, in target directed tapping tasks^47^, and in piano performance^48^. This type of behaviour, wherein participants constantly adapt to each other, is proposed to be the most energy efficient way of interpersonal coordination^19^.

Consistent with this, when analysing the lag coefficient of our participants we found that two out of three clusters showed patterns in line with mutual adaptation. These two clusters included the majority of our dyads and differed only in strength of the correlation while retaining the same pattern. However, we found a third cluster which exhibited a lag pattern not indicative of either leading-following nor mutual adaptation (see Figure 5b). The pattern, consisting of weak but positive correlations at all three lags, shows that there are little to no systematic interaction between the dyad members. Yet, the participants still achieve synchronized behaviour. A likely reason for this type of pattern to occur is if both participants are attempting to lead, and hence do not adapt to each other. This type of leading-leading-pattern is, to our knowledge, only previously observed in uncoupled interactions^4^.

The cluster found in our data did not show significant differences from the other two in terms of experience or synchronization measures, however, it did have a significantly higher occurrence of drummers and a higher occurrence of dyads consisting of drummers paired with drummers. In ensembles, and particularly in rhythmic bands, drummers are usually expected to be the main timekeeper. They usually produce rhythmic patterns consisting of multiple actions on differing metrical levels, as exemplified by the archetypical four-on-the-floor rhythm found in disco, pop, EDM, and rock. This pattern exists on a 4/4 meter, with the bass drum being hit every beat, the snare every other beat, and the hi-hat usually every half beat. To play such a rhythm a higher internal stability is needed than in for instance more melodic or monophonic instruments such as the voice, or the saxophone. Thus, one may argue that drummers are trained to be less metrically adaptive than other musicians.

It should, however, be noted that the cluster found in our data is relatively small, and individual differences or social factors could also play a large role in the observed data. For instance, synchronization is reduced when performing a repetitive task with a partner that arrives late^49^. As the two other clusters in our data also contained drummers, and dyads consisting of drummers paired with drummers, we would not generalize this effect to all drummers, but rather posit that this lag pattern may come about when two musicians that both are highly confident in their own internal RCM interact. To further explore the factors underlying if this type of interaction a follow-up study is necessary.

It is well known that musicians’ skills depend on instrument, level of expertise, and which genre of music they predominantly perform in^50–53^. On a neuronal level, these differences may manifest in low-level processing such as the mismatch negativity (MMN), a pre-attentive component of the auditory event-related potential considered to be a prediction error signal^54,55^. EEG studies have shown differences in sensitivity and amplitudes in MMN responses between musicians belonging to different genres^53,56^. On a behavioural level, drummers have been shown to be better at certain rhythmical tasks, yet the extent of these differences between types of musicians are not yet fully understood^57–59^. It might well be that these differences do not fully manifest in the common single-person experiments typically used in research on rhythm perception and production. Interacting with a computer-generated pacing signal, even if such signal is adaptive, does not fully capture the intricacies of joint action. Our study is one of the first to show that instrument-specific differences on an individual level may also impact synchronization strategies in interpersonal interaction.

## Conclusion

We tested the effect of dyads having a shared or non-shared rhythmic context model on synchronization in a joint finger tapping task. Having a non-shared model resulted in impaired synchronization at the start of a task, but the dyads recovered quickly. This suggests that in rhythmic joint action, musicians are able to efficiently adapt their own predictive models in order to facilitate interaction. Most dyads exhibited lag coefficient patterns indicative of mutual adaptation, except for a subset considering predominantly of drummers which showed a novel leading-leading-pattern suggesting little to no dyadic interaction. We find that a possible explanation for this is that the drummer’s role in normal musical performance may differ from many other instruments. Drummers often serve the role of timekeeper and motorically require a higher degree of internal synchronization of motion. The finding is of high interest, as it complements the strategies of leading-following or mutual adaptation in synchronization behaviour, by providing a third alternative wherein synchronization occurs without any apparent dyadic interaction. Further, it emphasises that differences between individual musicians, such as which instrument they play, also affect interpersonal synchronization strategies.

## Author Contributions

OAH contributed to the design, data collection, analysis, and interpretation of this work, as well as writing and revising the manuscript. IK was involved in the conception, design, analysis, and interpretation of the work, as well as revising the manuscript. MLK was involved in the interpretation of this work, as well as revising the manuscript. PV contributed to the conception, design, analysis and interpretation of this work, in addition to writing and revising the manuscript.

## Data Availability Statement

The data reported in this paper is available upon request. Due to data protection laws, a data processing agreement may be required.

## Acknowledgments

The authors thanks Suzi Ross, Frank Schulze, Marianne Tiihonen and Boris Kleber for assisting with data collection. OAH, MLK and PV are part of The Center for Music in the Brain, which is funded by the Danish National Research Foundation (DNRF117). IK is funded by the Danish Council for Independent Research – Technology and Production Sciences (0602-03001B).

